# Fitting high-resolution electron density maps from atomic models to solution scattering data

**DOI:** 10.1101/2023.06.02.543451

**Authors:** Sarah R. Chamberlain, Stephen Moore, Thomas D. Grant

## Abstract

Solution scattering techniques, such as small and wide-angle X-ray scattering (SWAXS), provide valuable insights into the structure and dynamics of biological macromolecules in solution. In this study, we present an approach to accurately predict solution X-ray scattering profiles at wide angles from atomic models by generating high-resolution electron density maps. Our method accounts for the excluded volume of bulk solvent by calculating unique adjusted atomic volumes directly from the atomic coordinates. This approach eliminates the need for a free fitting parameter commonly used in existing algorithms, resulting in improved accuracy of the calculated SWAXS profile. An implicit model of the hydration shell is generated which uses the form factor of water. Two parameters, namely the bulk solvent density and the mean hydration shell contrast, are adjusted to best fit the data. Results using eight publicly available SWAXS profiles show high quality fits to the data. In each case, the optimized parameter values show small adjustments demonstrating that the default values are close to the true solution. Disabling parameter optimization results in a significant improvement of the calculated scattering profiles compared to the leading software. The algorithm is computationally efficient, showing more than tenfold reduction in execution time compared to the leading software. The algorithm is encoded in a command line script called denss.pdb2mrc.py and is available open source as part of the DENSS v1.7.0 software package (https://github.com/tdgrant1/denss). In addition to improving the ability to compare atomic models to experimental SWAXS data, these developments pave the way for increasing the accuracy of modeling algorithms utilizing SWAXS data while decreasing the risk of overfitting.

**Statement of Significance:** Accurate calculation of small and wide-angle scattering (SWAXS) profiles from atomic models is useful for studying the solution state and conformational dynamics of biological macromolecules in solution. Here we present a new approach to calculating SWAXS profiles from atomic models using high resolution real space density maps. This approach includes novel calculations of solvent contributions that remove a significant fitting parameter. The algorithm is tested on multiple high quality experimental SWAXS datasets, showing improved accuracy compared to leading software. The algorithm is computationally efficient and robust to overfitting, paving the way for increasing the accuracy and resolution of modeling algorithms utilizing experimental SWAXS data.

## 1. Introduction

Solution scattering provides information on the structure and dynamics of biological macromolecules in aqueous solutions, yielding complementary information to high-resolution structural techniques such as X-ray crystallography, cryo-electron microscopy (cryoEM), and nuclear magnetic resonance (NMR) (1-5). Ab initio modeling approaches have enabled the determination of low-resolution 3D shapes and density maps directly from 1D small angle X-ray scattering (SAXS) profiles (6-9). We have previously described our approach for low-resolution ab initio generation of density maps from X-ray or neutron scattering data, called DENSS (9). DENSS operates by iterating between the real space and reciprocal space domains using the Fast Fourier Transform (FFT) (10), enforcing appropriate restraints in each domain (i.e., 1D SAXS data in reciprocal space and 3D solvent flattening in real space). One advantage of using density to describe the object is the potential ability to model high-resolution fluctuations in density that may reflect features in the wide angle X-ray scattering (WAXS) regime (11).

Hybrid modeling approaches combining high-resolution atomic models with solution scattering data have proven useful for improving the resolution, accuracy, and interpretability of both small and wide angle X-ray scattering data (SWAXS) (12-15). Generating hybrid models requires the accurate calculation of SWAXS profiles from atomic models accounting not only for the macromolecule of interest, but also solvent contributions, including both the displaced bulk solvent (i.e., the “excluded volume”) and the hydration shell surrounding the surface of the macromolecule (16). Many convenient algorithms have been proposed for fitting such solvent terms to SWAXS profiles, such as CRYSOL, FoXS, and others (16-19), with CRYSOL being the most widely used. These algorithms most often include fitting parameters to adjust for differences in solvation or solution conditions, or for experiment errors. Several more advanced and computationally expensive algorithms exist for modeling solvent terms explicitly, including utilizing molecular dynamics (MD) simulations, to fit high resolution WAXS data accurately with minimal free parameters (20-23).

We sought to calculate a high-resolution density map from an atomic model that matches an experimental SWAXS profile to wide angles. Existing approaches calculate scattering profiles in reciprocal space directly from atomic model coordinates. These approaches typically use the Debye equation (17,24), spherical harmonics approximations (16), or 3D Fourier transforms (18,22,23). The Debye equation requires a double summation over every pair of atoms and therefore the computational cost scales as the square of the number of atoms. For larger particles with many atoms, this cost is often a limiting factor when performing modeling requiring repeated calculations from potentially thousands of candidate models. To deal with this computational cost, alternative approaches have been developed, including using coarse graining to approximate the particle with fewer atomic groups (for example, replacing all the atoms of a protein residue with a specially modified form factor representing the entire residue), resulting in significant speed improvements with acceptable accuracy in the small angle regime (17,25-27). Others have sped up the calculation by using a spherical harmonics expansion to approximate the scattering profile rather than the Debye equation (16). Another method is the “cube” method (28) that several algorithms employ which calculates an explicit 3D Fourier transform from atomic coordinates using a cubic lattice, and then performs spherical averaging numerically in reciprocal space using a sufficient number of particle orientations to generate the 1D scattering profile (18,20-23).

Many approaches directly calculate the scattering profile in reciprocal space from the atomic model, with no intermediate real space density. However, as our aim is to generate an accurate high-resolution density map matching the experimental 1D SWAXS profile, we require calculating a density map in real space directly from the atomic coordinates. We briefly review the general theory behind these calculations below before describing our approach to calculate the real space high-resolution density of the macromolecule from an atomic model including the excluded volume and hydration shell which results in high quality fits to experimental SWAXS data. Our approach includes a unique method for estimating the volumes of atoms for the excluded volume calculation, resulting in the removal of a free fitting parameter. We show that the best fits obtained result in narrow distributions near the default values for the two remaining fitting parameters (namely bulk solvent density and hydration shell contrast). The method is flexible, including the ability to evaluate different methods of excluded volume and hydration shell contrast, easy to use, and faster than the leading software. We show that our approach is capable of generating high-resolution density maps that accurately fit experimental data from publicly available SWAXS datasets. In addition to providing a new tool for fitting atomic models to SWAXS profiles out to wide angles, this approach provides the foundation for the future development of tools enabling hybrid density-based modeling, which promises to significantly increase the accuracy and resolution of density reconstructions from SWAXS data.

## 2. Methods

To begin, there are three significant components of particle scattering that all contribute to the final 1D solution scattering profile. Equation 1 describes these three terms, the contribution from the particle atoms in vacuum and two additional solvent contributions including the excluded volume and the hydration shell:

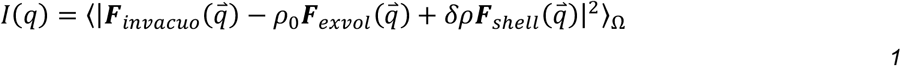

where *I*(*q*) is the 1D scattering profile intensity as a function of momentum transfer *q*, where 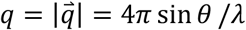 in inverse angstroms (Å^-1^), 2*θ* is the scattering angle, and *λ* is the wavelength of the incident radiation, ***F***_*invacuo*_ is the 3D complex structure factor in reciprocal space in vacuum (***F*** is sometimes called “scattering amplitudes” in other works), *ρ*_0_ is the average bulk solvent density, ***F***_*exvol*_ is the structure factor corresponding to the excluded volume of bulk solvent displaced by the solute, *δρ* is the contrast of the hydration shell, ***F***_*shell*_ is the structure factor corresponding to the hydration shell density, and ⟨ ⟩_Ω_ refers to the spherical average over all 3D orientations to obtain the 1D intensity profile.

The *in vacuo* calculation requires the atomic form factor to calculate the *in vacuo* structure factor, ***F***_*invacuo*_, as shown in Equation 2.

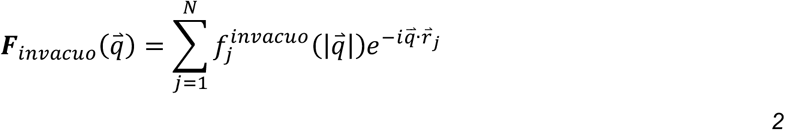

where 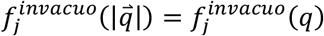 is the spherically symmetric atomic form factor as a function of the modulus of the scattering vector 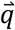, and the summation is over all *N* atoms. The atomic form factor is most often calculated by parameterizing the atomic scattering as a sum of four Gaussian functions and a constant offset using the nine Cromer-Mann coefficients (two per Gaussian plus one offset) (Equation 3), where each atom type has a unique set of coefficients (29). This formulism is described by Equation 3

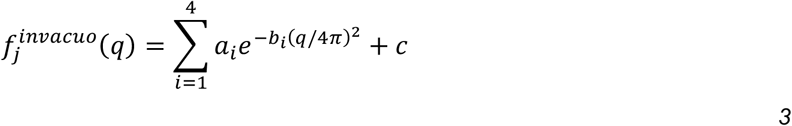

where *a*_*i*_, *b*_*i*_, and *c* are the nine Cromer-Mann coefficients for the atomic form factors tabulated in the International Tables of Crystallography (30).

The second term in Equation 1, ***F***_*exvol*_, refers to the scattering from the bulk solvent that has been displaced by the solute, often called the “excluded volume”. The difference between the *in vacuo* density and the excluded volume gives rise to what is commonly called the “contrast” of the particle in the solvent. Since the bulk solvent is largely disordered, it is most often considered as a flat uniform density with an average value of *ρ*_0_. A typical value for *ρ*_0_ is that of pure water at room temperature, 0.334 e^-^/Å^3^, but this value can be affected by temperature or other components in the aqueous solution including salts and buffer molecules that can alter the average bulk density value. Some approaches model the excluded volume term as a simple flat value inside the envelope defined by the solute (18,20-23). Other methods use molecular dynamics simulations to model the solute in explicit solvent, and separately model the bulk solvent, taking the difference of the two calculations as the contrast (21,22,31). Several of the more popular methods such as CRYSOL and FoXS model the excluded volume as a simple isotropic Gaussian centered at the coordinates of the atomic model of the solute, an approach originally described by Fraser, et. al. (16,17,32). This formulation can be expressed by Equation 4

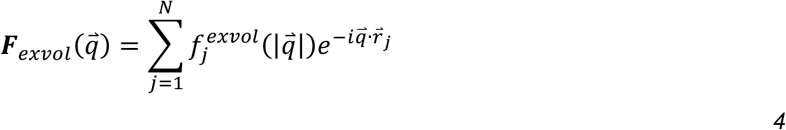

where

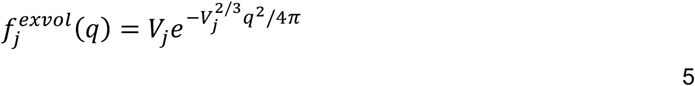

where *V*_*j*_ is the volume of atom *j*, which is calculated assuming a hard sphere representation of the atom as 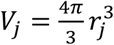, where *r*_*j*_ is the radius of the atom. It is important to note here that the radius of the atom is not trivially obtained. The goal of the excluded volume estimation is to approximate the total volume of bulk solvent excluded by the macromolecule. Thus, the radius of each atom plays a critical role in the total volume estimation, and small adjustments to the atomic radii have significant impact on the shape of the SWAXS curve. Since neighboring atoms in a macromolecule such as a protein are covalently bound, much of the electron density participating in bonding overlaps between two atoms. This results in the total excluded volume being less than the sum of the individual volumes based on the van der Waals radii of the atoms (18), such as those given by Bondi (33). This gives rise to “observed” volumes listed by Fraser, et. al. (32), and which are used by programs such as CRYSOL. It is noteworthy that multiple values for calculated and observed atomic volumes are reported by Fraser, et. al., demonstrating that these values are in fact not well defined in the literature (18,19). For this reason, programs such as CRYSOL and FoXS include an adjustable fitting parameter called the global expansion factor that effectively increases or decreases the average atomic radius to best fit the experimental data. The common approach of describing each atom of a particular type as having a single radius (and thus volume) does not consider the variations in bond lengths and in the number of overlapping atoms throughout the molecule.

The third and final term in Equation 1, ***F***_*shell*_, arises from the observation that solvent molecules in the immediate vicinity of the surface of a macromolecule are slightly more ordered on average than in the bulk resulting in a slightly higher average density near the surface (34). This region is referred to as the hydration shell. Several programs such as CRYSOL model the hydration shell as an implicit layer of uniform contrast relative to the bulk, with a thickness of 3 Å (about the diameter of a water molecule), and a contrast of about 0.03 e^-^/Å^3^, which is adjusted to best fit the experimental data. While simple, the implicit shell model has proven effective at approximating the hydration shell and fitting experimental SAXS profiles. Several alternative approaches have also been proposed that employ explicit solvent models. These are often computationally expensive approaches which require molecular dynamics simulations or sophisticated models of the hydration shell (20-23,31,35). However, the major advantage of these explicit solvent methods is the reduction of free fitting parameters, enabling accurate SWAXS calculations even at wide angles with few or no free parameters.

We describe below our approach to generating a high-resolution density map from an atomic model that has a corresponding scattering profile that matches the 1D experimental SWAXS pattern to wide angles. Our desire is the calculation of real space density from an atomic model which can be used with tools such as DENSS to model SWAXS data. This requires the ability to represent the object as a density map and apply a forward and inverse FFT (10) to switch between the real space and reciprocal space domains. To do so, we first calculate density *ρ* and then perform the Fourier transform (FFT[*ρ*]) to obtain the corresponding ***F*** in Equation 1 (as opposed to directly calculating the ***F***’s in reciprocal space).

### 2.1. Real space density calculation

To begin, we set up a real space density map as a cubic grid of voxels, similar to the setup of *ab initio* density reconstruction in DENSS. In this case, we are particularly interested in high-resolution density maps, and therefore we set the default voxel size to 1 Å to provide sufficient sampling of the atomic model and solvent terms. The size of the box is then dictated by the size of the particle, with the default box size set to three times the maximum dimension of the particle (D_max_), to more than satisfy the minimum box size required by Shannon sampling of 2*D_max_ (36-38). This approach results in more finely sampled maps than conventional DENSS, which requires greater memory. If the number of voxels using the default settings exceeds 256 on a side, then the voxel size is increased to accommodate this, resulting in decreased sampling in real space that may affect very wide scattering angles (since *q*_*max*_ = 2*π*/*dx*, where *dx* is the voxel spacing), though such wide scattering angles are rarely collected in SWAXS experiments. Following calculation of the real space density map, the FFT is used to calculate the structure factors in reciprocal space. Finally, as in DENSS, the 3D intensities are calculated, and the spherically averaged 1D SWAXS profile is calculated by binning intensities according to their *q* value. Further details for this calculation are given below.

### 2.2. Real space in vacuo density calculation

To calculate *ρ*_*invacuo*_ we use the corresponding real space form of Equation 3 which uses a four Gaussian sum to represent the atomic form factor defined by the International Tables of Crystallography (30). Additionally, we account for a possible atomic B-factor for each atom. We use a radial Fourier transform to obtain Equation 6

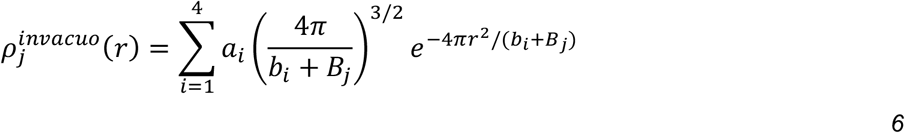

where *r* is the distance from the atomic coordinate to the grid point and *B*_*j*_ is the atomic B-factor. Note that we have removed the offset *c* in this equation as this results in a delta function in real space at *r* equal to zero and subsequently reparametrized *a*_*i*_ and *b*_*i*_. Since the density quickly decays as *r* increases, for speed the density is only calculated for voxels within 3 Å of the atom center (or an appropriately larger region if B-factor is included). To ensure the correct number of electrons is accounted for, after the density is calculated for an atom, the density is scaled such that the total number of electrons equals the expected number of electrons for the atom type.

Assuming a B-factor of zero for atoms results in density very closely situated near the center of the atom using Equation 6. In some cases, such as for nitrogen, a significant fraction of the density is less than 0.03 Å from the atom center. Since the default voxel size is 1 Å this often results in the central spike of density not being sufficiently sampled by the grid. To adjust for this, a small B-factor (∼7 Å^2^, equivalent to a 0.3 Å atomic displacement) is added to all atoms to spread out the density sufficiently to nearby voxels. This improves the real space sampling while having a modest effect on the reciprocal space scattering calculation. To correct for this modest effect, B-factor sharpening is performed (i.e., using -7 Å^2^) in 3D reciprocal space prior to calculating the 1D SWAXS profile. This B-factor correction to deal with voxel sampling issues is in addition to any B-factor from the atomic model, which by default is set to zero (explained in more detail below).

### 2.3. Excluded volume

To calculate the density corresponding to the excluded volume, we follow the approach of Fraser, et. al., (and implemented in CRYSOL, FoXS and others) to model the excluded volume of each atom as an isotropic Gaussian sphere centered at each solute atom, whose volume is equal to the volume of the solute atom, as described by Equation 7

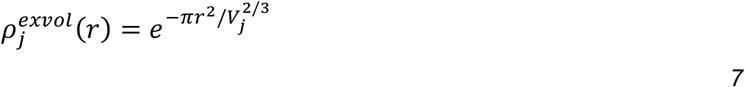

It is important to note that the volume of each atom, *V*_*j*_, is not simply the van der Waals volume of the solute atom. Due to atomic bonding, neighboring atoms significantly overlap one another, and this overlap must be accounted for when estimating the total volume of bulk solvent displaced by the macromolecule, resulting in an “adjusted” volume for each atom. A common method for estimating the adjusted volume of the solute atom in programs such as CRYSOL, FoXS, Pepsi-SAXS, and others (16,17,19) is to assign the same atomic volume to each atom of a particular type (e.g., hydrogen, carbon, etc., or more generally each atomic group including hydrogens), where the adjusted volumes are based on previously reported values (e.g., values reported in Fraser, et. al.). This is typically followed by fitting a global expansion factor which effectively increases or decreases the volume of all atoms by a fraction (up to ∼30% in CRYSOL) to best fit the experimental data to adjust for errors in the volume estimation.

To avoid this additional fitting parameter, we developed a method for estimating the volume of an atom that is unique to each individual atom based on distance calculations between neighboring atoms in the atomic model and thus to account for variations in bonding and local environment. To calculate these unique, per-atom volumes, each atom is isolated and placed into its own miniature Boolean voxel grid with very fine sampling (n=16 voxels per side, or about 0.2 Å voxels, depending on the van der Waals radius). Each voxel is set to either True or False based on whether or not the voxel falls within the boundaries of the primary atom while accounting for overlapping atoms. First, all voxels within the van der Waals radius of the primary atom center are set to True, and the rest to False (Fig. 1). Second, all atoms are selected that are within 5 Å of the primary atom. Third, for each of the selected overlapping atoms, the equation of the plane forming the sphere-sphere intersection of the pair of atoms is calculated. Voxels that are further away from the primary atom center than this plane, i.e., voxels that are on the “other side” of the plane, are considered to be a part of the neighboring atom and are thus set to False (Fig. 1). This calculation is repeated for all overlapping atoms. Finally, the total volume of the remaining voxels is calculated and used in Equation 7 as the adjusted volume for that atom. This approach allows for more precise per-atom volume estimations as each individual atom is assigned a volume based on the volume of space it takes up while accounting for nearby bonded atoms and is based on the observed (or at least restrained) bond lengths calculated from the experimentally determined high-resolution atomic model.

**Figure 1.**
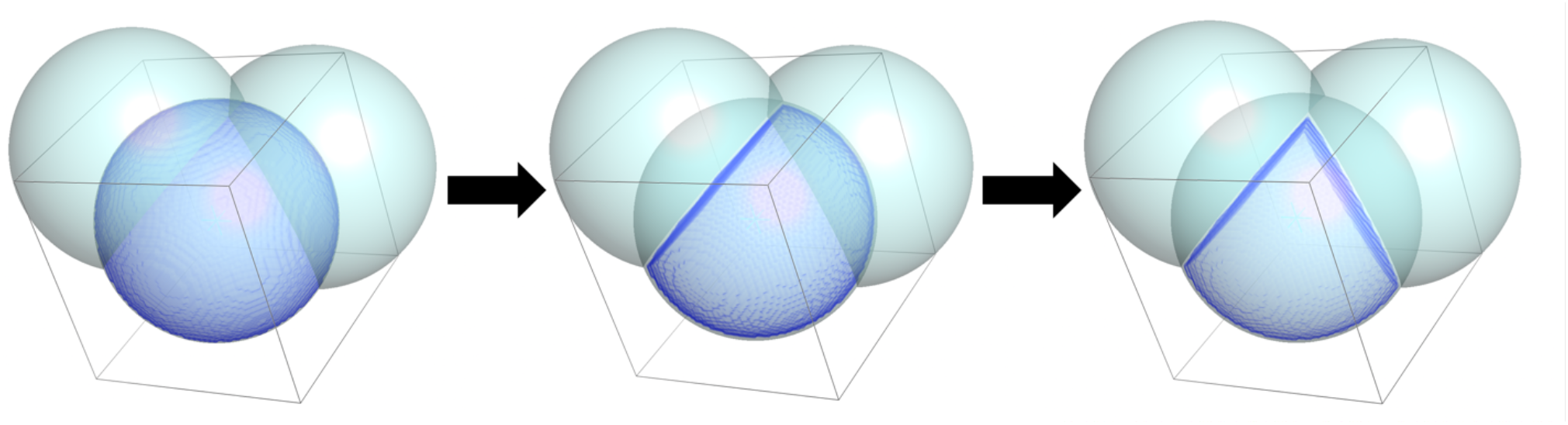
Schematic showing the calculation of unique “adjusted” volumes of solute atoms. Three carbon atoms are shown as a cyan sphere, each with a van der Waals radius of 1.7 Å. The central atom is overlaid with a blue sphere representing the volume of the atom that does not overlap the neighboring carbon atoms. From left to right, the volume of the central atom overlapping with the neighboring carbon atoms is removed, resulting in the spherical wedge on the right. The volume of the spherical wedge is taken as the adjusted volume of the central atom and used in Equation 7 for the calculation of the excluded volume density map.

The calculation of unique volumes for each atom can take several minutes for large macromolecules. To improve the computational efficiency of this step, we generated a dictionary of average adjusted atomic volumes for atomic groups found in amino acids and nucleic acids. The program Reduce (39) as implemented in the Phenix package (40) was used to add hydrogens to ∼2000 atomic models downloaded from the Protein Data Bank (PDB) (41). The per-atom unique volume calculation described above was then used to calculate the adjusted atomic volume for each atom name (e.g., *N, C*_*α*_, *C, O, C*_*β*_, *H*_*α*_, *H*_*β*2_, etc.) for each chemical group found (e.g., amino acids, nucleic acids, etc.). For each particular atom name, the adjusted volumes for all instances of that atom name were calculated, averaged, and entered into the dictionary. When performing the SWAXS calculation for a new model, this dictionary is used to look up the adjusted volume for each atom, rather than calculating the adjusted volumes directly from the atomic model. For any atoms in the model not found in the dictionary, the unique volumes are calculated explicitly. Using this dictionary as a look up table for adjusted volumes significantly decreases the run time (though the option to explicitly calculate them for each model is available). Fig. 2 shows the distributions of volumes for different amino acids for a set of eight example atom names. Note that the distributions are narrow for each amino acid and atom name, showing that the mean volume is a suitable representation for the atomic volume of a particular atom in an atomic model.

**Figure 2.**
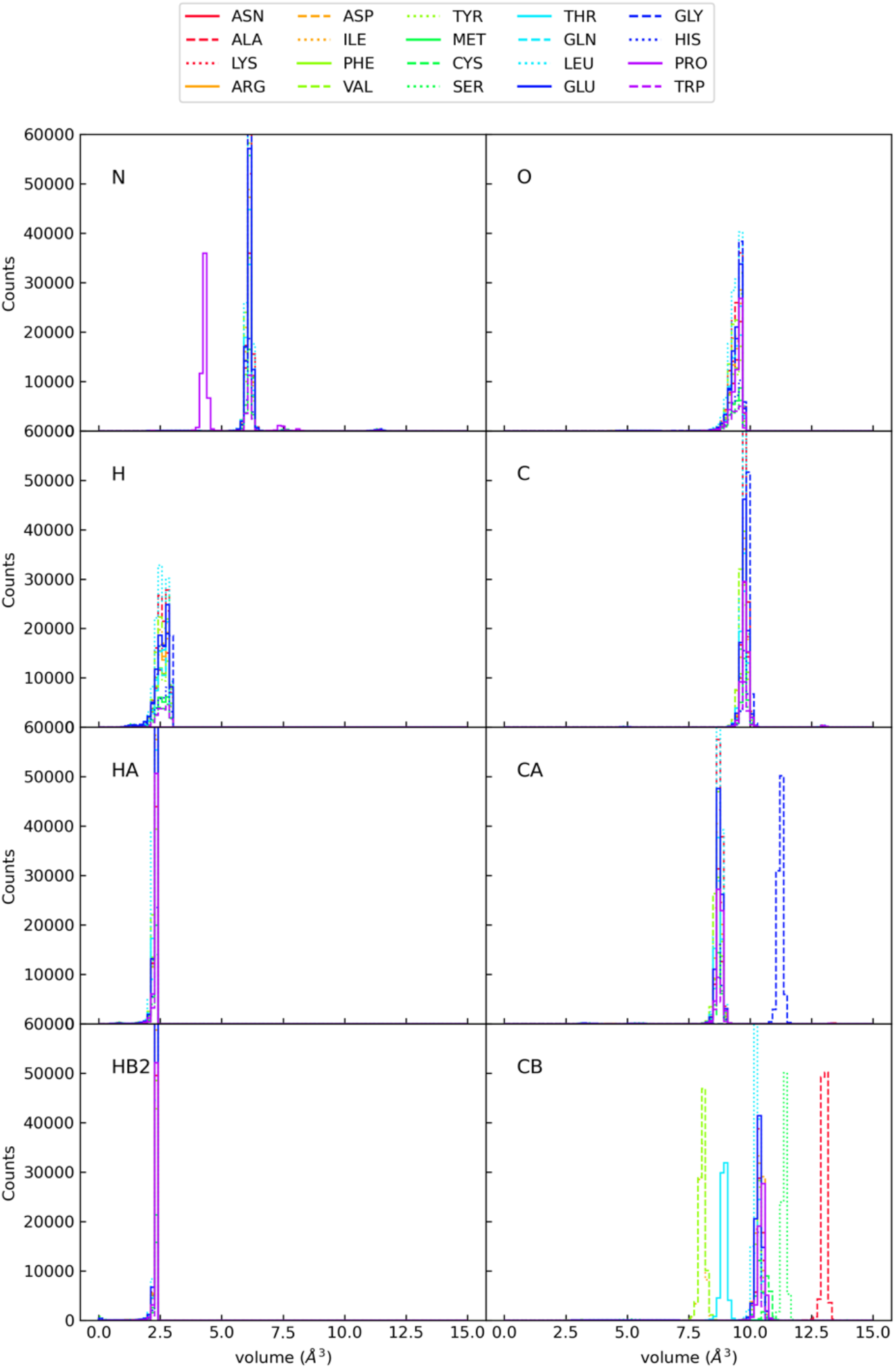
Distributions of atomic volumes calculated for a selection of specific atoms for all twenty amino acids. Volumes were calculated using the approach described in the text for a set of nearly 2000 PDB files. The mean value for each atom name is recorded and used as the adjusted atomic volume in denss.pdb2mrc.py.

However, the mean volume often differs for different amino acids for the same atom type and even for the same atom name, demonstrating that independent volume assignments for each atom name, including distinguishing between amino acids (or other groups), is required for accurate estimation of the adjusted volume of an atom. This shows that using a single global atom radius for all atoms of a particular type (i.e., hydrogen, carbon, nitrogen, etc.) is not discriminatory enough to accurately estimate the excluded volume of a particle, but that discriminating at the level of specific atom names for different amino acids is sufficient. This increased accuracy in estimating excluded volume is key to enabling the removal of the global expansion factor as a fitting parameter.

This approach is likely to underestimate the volume of the excluded bulk solvent, since it does not account for small voids between atoms that solvent molecules cannot reach, the so called solvent excluded surface. To account for this, scale factors were determined to correct the volume of each atom type (for hydrogen, carbon, nitrogen and oxygen). To determine these scale factors, a benchmark SWAXS profile was downloaded from the SASBDB archive (42) along with the corresponding high-resolution atomic model for lysozyme (SASDCK8). The scale factors that resulted in the best fit to the experimental data (using the fitting routine described below, while restraining *ρ*_0_ to be 0.334 e-/Å^3^ to avoid overfitting) were recorded and tested with several other datasets for validation (see Results section). As shown in the Results section, despite obtaining these scale factors from only one high quality SWAXS profile, they were found to be suitable for all other SWAXS profiles tested. The advantage of this approach over others is the removal of one of the free fitting parameters, namely the global volume expansion factor, as the unique, per-atom excluded volumes are now calculated directly from the atomic model.

### 2.4. Hydration shell

The third and final term is density corresponding to the hydration shell. The water molecules that associate with the surface of the macromolecule form layers, or shells, whose peak is located approximately one water molecule radius from the macromolecule surface. The amount of ordering and thus level of contrast varies throughout the surface based on how tightly water molecules associate with that region of the surface, often based on the polarity of the region. In addition to using MD, programs such as HyPred, AquaSAXS, 3D-RISM and others (20,22,23) have generated sophisticated methods of modeling this variation in contrast around the particle surface without requiring full MD simulations for each macromolecule. Simpler implicit hydration shell models, available in programs such as CRYSOL, FoXS, and others (16-19), assume a uniform density throughout the shell, followed by fitting of the contrast value using experimental data. We sought to develop an implicit shell model for simplicity and to enable fitting of the contrast, but also to develop a model that would allow the capture of the variation in the shell contrast as a function of distance from the solute surface and to perform this calculation in real space as a density map.

To begin, the center of the water shell is set to the macromolecular surface (defined by the van der Waals radii of the surface atoms) plus the radius of a water molecule (set to 1.4 Å). Rather than modeling the shell as a uniform contrast of a set thickness, the shell is modeled using the real space form factor of a water molecule. A distance transform is calculated to determine the distance between each voxel in the shell and the center of the water shell. Next, the density of the shell is calculated for each voxel around the surface using the distance transform with the real space form factor of a water molecule.

While this approach generates a real space density map of the contrast of a hydration shell, it implicitly assumes that the water is rigid at the surface. As explained above, in the case of a hydration shell, water molecules in the shell are regularly exchanging with the bulk and are far less ordered on average than a single rigid water molecule. This disorder will affect the overall scale of the contrast. To allow for variations in the hydration shell for different macromolecules, the mean contrast of the shell is adjusted to best fit the data (the fitting procedure is described below). However, to estimate a reasonable default value, the mean contrast of the shell is set to 0.019 e^-^/Å^3^, which is the average value found for the eight datasets described below. An example of a shell for PDB 6LYZ (43) is shown in Fig. 3. The advantage of this approach is that it combines the simplicity of implicit shell models with the ability to model varying contrast as a function of distance from the solute surface (in contrast to a simple uniform shell). This approach allows for the water shell to penetrate the interior cavities and channels of the molecule (31) (Fig. 3); however, it does not currently attempt to model variations in contrast across the surface of the macromolecule as possible with more sophisticated and computationally expensive methods (20-22,31,44,45).

**Figure 3.**
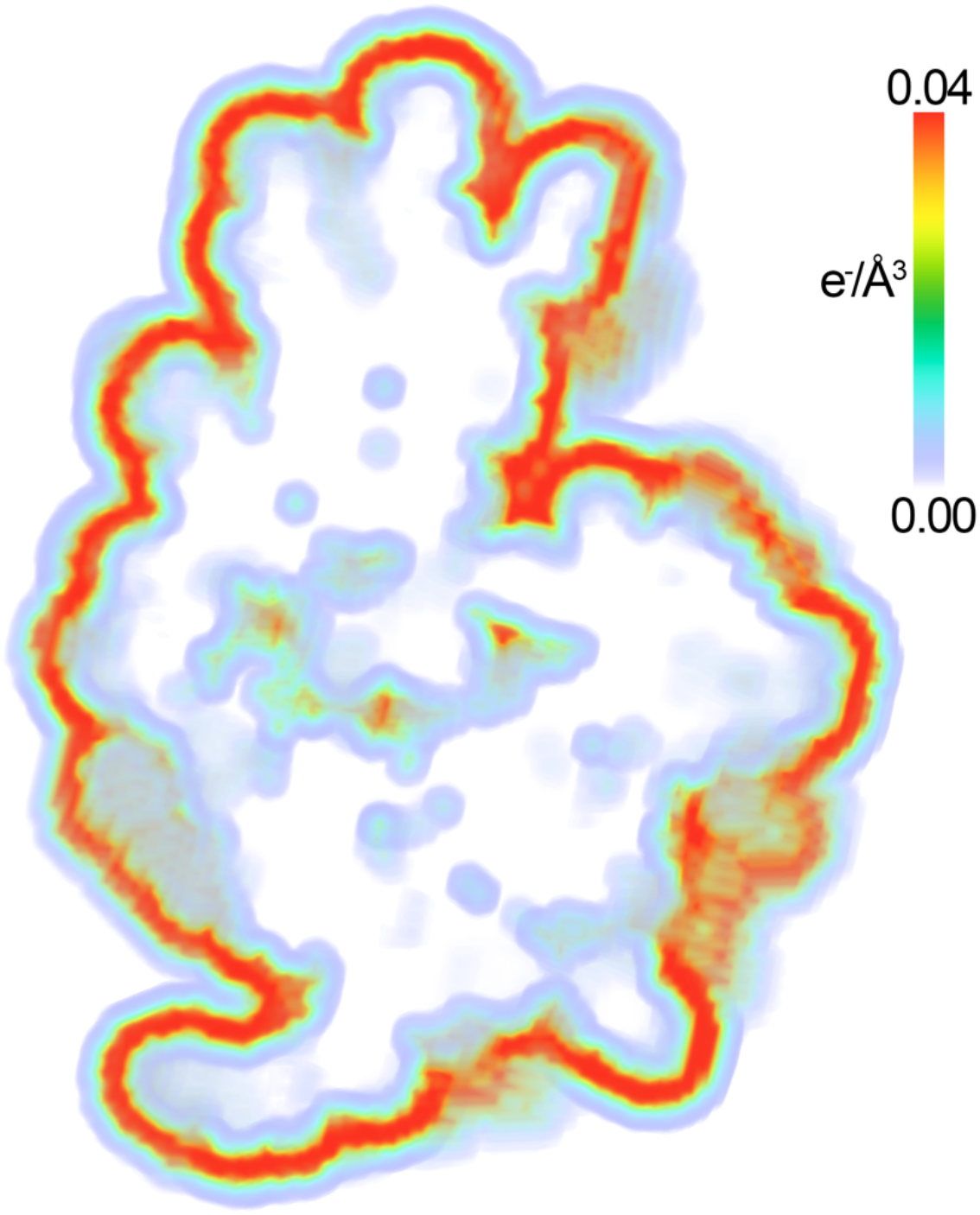
Electron density map of the hydration shell of lysozyme. The electron density of the hydration shell is shown as contrast relative to the bulk solvent and is colored in units of e^-^/Å^3^ as shown in the color bar using the “volume” representation in PyMOL (46).

### 2.5. Fitting/minimization procedure

Once the real space density maps are calculated for the *in vacuo* macromolecule, the excluded volume, and the hydration shell, the total scattering profile is calculated using fast Fourier transforms (FFT) (10) and Equation 1. The structure factors of the three density maps are calculated using the FFT. For the *in vacuo* structure factors, ***F***_*invacuo*_, the B-factor sharpening is additionally performed in reciprocal space as described above prior to summation of the three terms. The bulk solvent density, *ρ*_0_, and the hydration shell contrast, *δρ*, in Equation 1 are then optimized in reciprocal space to best match the experimental data.

A numerical optimization routine was implemented to enable fitting of the calculated density to experimental SWAXS data. Experimental scattering profiles are often highly oversampled, in particular when compared to the reciprocal space sampling resulting from the FFT. The default box size of three times the particle maximum dimension results in approximately 1.5 sampled points in the calculated scattering profile for every Shannon channel (9,36-38), which are not sampled at the same *q* values as the experimental data. To enable comparison of the coarsely sampled calculated scattering profile with the finely sampled experimental scattering profile, the calculated scattering profile is interpolated to the *q* values in the experimental profile using a third order cubic spline. The *χ*^2^ between the calculated intensities and the experimental intensities is then calculated as a measure of the goodness of fit according to Equation 8

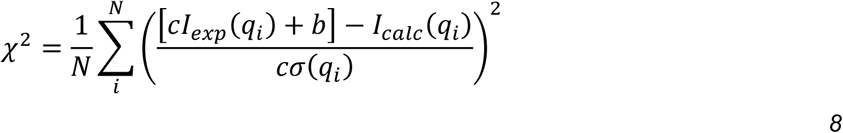

where *N* is the number of experimental data points *i* in the data, *I*_*exp*_ is the experimental SWAXS profile measured at *q*_*i*_ values with experimental errors *σ, I*_*calc*_ is the calculated and interpolated scattering profile, *c* is a constant scale factor determined by least squares, and *b* is an optional offset to correct for errors in background subtraction also determined by least squares (disabled by default). Numerical optimization using the Nelder-Mead simplex algorithm (47,48) is used to obtain the best values for *ρ*_0_ and *δρ*.

## 3. Results

The approach outlined above has been implemented in a command line script called denss.pdb2mrc.py as part of DENSS v1.7.0 (9). To validate the accuracy of denss.pdb2mrc.py using experimental data, eight SWAXS datasets from the benchmark section of the SASBDB (42) online database were downloaded along with the corresponding atomic models. Reduce (39) was used to add hydrogens to the atomic models. Most datasets have a maximum measured *q* value of 1.79 Å^-1^, well into the WAXS regime, suitable for testing the accuracy of the algorithm out to wide angles. A wide range of particle sizes are available in this dataset, which includes cytochrome C (11.7 kDa, SASDCH8), lysozyme (14.3 kDa, SASDCK8), ribonuclease A (16.5 kDa, SASDAN2), myoglobin (17.1 kDa, SASDAK2), carbonic anhydrase (29.0 kDa, SASDCG8), bovine serum albumin (BSA) (66.2 kDa, SASDCF8), glucose isomerase (172.4 kDa, SASDCJ8), beta amylase (224.0 kDa, SASDCE8), and apoferritin (475.2 kDa, SASDCD8). Results of the fits shown for each SWAXS profile using denss.pdb2mrc.py with default parameters are shown in Fig. 4. Fits using CRYSOL (from ATSAS v3.1.3, (16,49)) with the same atomic model (using 50 harmonics, explicit hydrogens, and 256 points) are also shown for comparison. Comparison of *χ*^2^ values (Table 1) shows that the method implemented in denss.pdb2mrc.py is as accurate as CRYSOL (i.e., *χ*^2^ is lower than CRYSOL for some datasets and higher for others). These results confirm that our approach generates accurate electron density maps from atomic models that fit experimental SWAXS profiles out to wide angles, despite using only two free fitting parameters.

**Table 1.** Parameters for fitting atomic models to SWAXS data. ID is the SASBDB ID for each sample, N_atoms_ is the number of atoms (including hydrogens) in the model, ρ_0_ is the bulk solvent density, δρ is the mean contrast of the hydration shell, 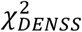 is the goodness of fit of the scattering profile calculated by denss.pdb2mrc.py using Equation 8, 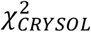 is the goodness of fit of the scattering profile by CRYSOL evaluated using Equation 8 (Note: for fair comparison, these values were recalculated using the denss.pdb2mrc.py code with the fits output by CRYSOL, though this resulted in virtually identical values to those reported by CRYSOL).

| ID | Sample | $N_{\text{atoms}}$ | $\rho_0$ ($e^-/\text{\AA}^3$) | $\delta\rho$ ($e^-/\text{\AA}^3$) | $\chi^2_{\text{DENSS}}$ | $\chi^2_{\text{CRY SOL}}$ |
| --- | --- | --- | --- | --- | --- | --- |
| SASDCH8 | cytochrome C | 1744 | 0.326 | 0.014 | 24.35 | 20.12 |
| SASDAN2 | ribonuclease | 1852 | 0.341 | 0.038 | 2.46 | 2.86 |
| SASDCK8 | lysozyme | 1959 | 0.334 | 0.019 | 1.76 | 2.08 |
| SASDCG8 | carbonic anhydrase | 4035 | 0.337 | 0.023 | 18.30 | 21.44 |
| SASDCF8 | BSA | 8387 | 0.337 | 0.018 | 14.98 | 11.34 |
| SASDCJ8 | glucose isomerase | 23848 | 0.335 | 0.025 | 13.69 | 9.73 |
| SASDCE8 | beta amylase | 30708 | 0.336 | 0.002 | 50.65 | 54.15 |
| SASDCD8 | apoferritin | 66456 | 0.341 | 0.023 | 91.00 | 353.18 |

**Figure 4.**
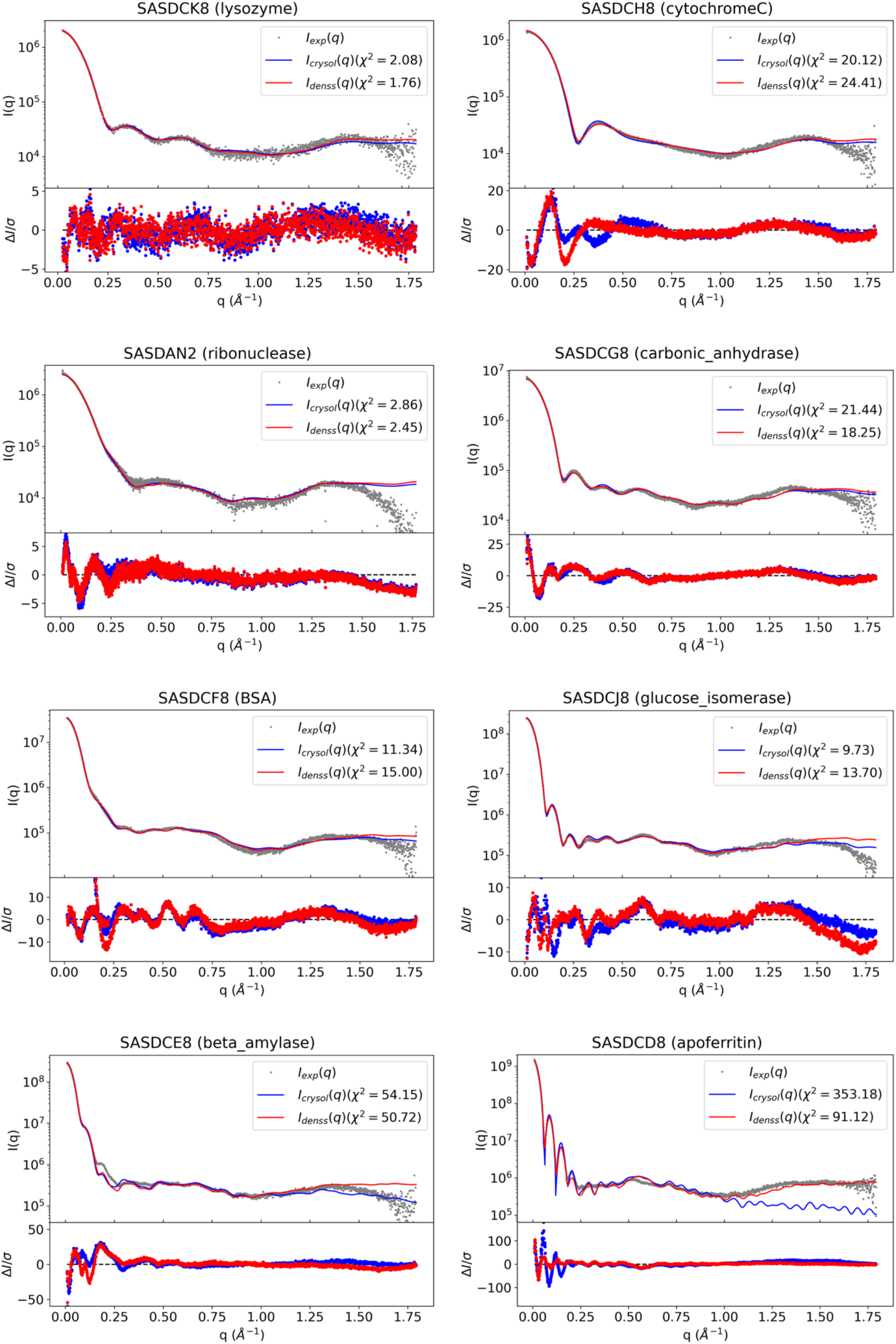
Fits and residuals of calculated scattering profiles from atomic models compared to experimental SWAXS data using denss.pdb2mrc.py and CRYSOL. For each sample, the top plot shows the experimental scattering data from the SASBDB entry as gray dots, and the fits to the data using CRYSOL (blue solid line) and denss.pdb2mrc.py (red solid line). The bottom plot shows the normalized residuals using the same color scheme, which are calculated as ΔI/σ, where ΔI is the difference between the data and the fit, and σ is the experimental error. The χ^2^ is shown in the legend for each sample for both methods.

Fig. 5 shows an example of the calculated density map for the *in vacuo* density, the excluded volume density, and the hydration shell density for lysozyme (SASDCK8), along with the final density. Table 1 shows the fitting parameters for denss.pdb2mrc.py (including the bulk solvent density and the hydration shell contrast) for each dataset along with the quality of the fit. The fitted values for both *ρ*_0_ and *δρ* are tightly distributed near the default values (mean *ρ*_0_ = 0.336 ± 0.004, mean *δρ* = 0.019 ± 0.009), showing that the calculation makes minimal adjustments to the two fitting parameters (defaults of 0.334 and 0.019, respectively).

**Figure 5.**
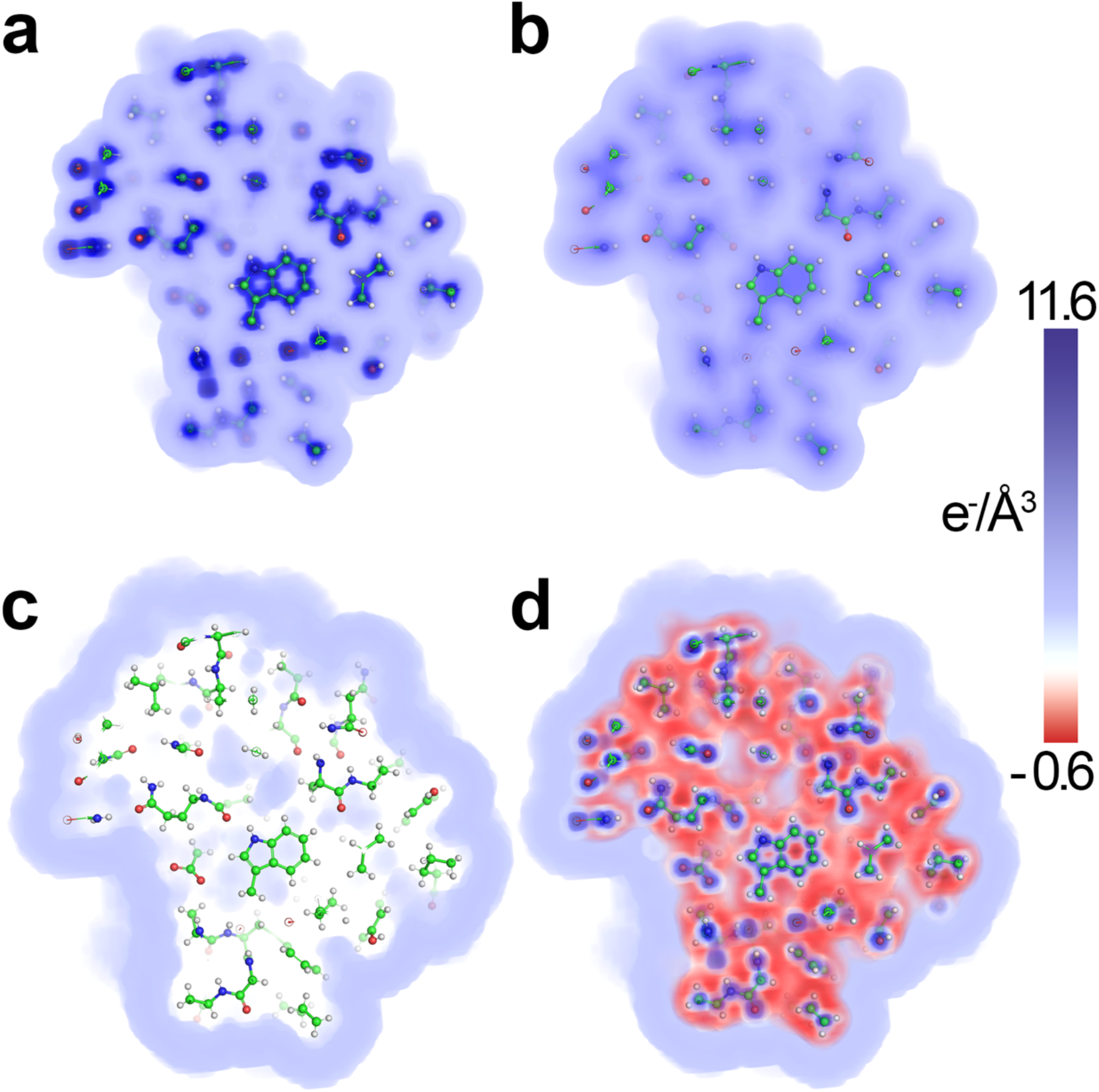
Electron density maps of lysozyme calculated from atomic models using experimental SWAXS data. Electron density maps were calculated from the atomic model of lysozyme deposited in the SASBDB entry SASDCK8 and fitted to the experimental SWAXS data using denss.pdb2mrc.py. Density maps shown are cross sections and include (a) in vacuo, (b) excluded volume, (c) hydration shell, and (d) total (i.e., in vacuo – excluded volume + hydration shell). The density maps are colored according to density shown in the color bar in units of e^-^/Å^3^ using PyMOL (46).

The observation that the distributions for the two fitting parameters, *ρ*_0_ and *δρ*, were narrowly centered on the default values motivated us to test the effect of disabling fitting entirely. If the default values return high quality fits without optimization, that would suggest that it may be possible to avoid fitting, which would remove a significant barrier to modeling algorithms using SWAXS data, since the lack of information in SWAXS profiles may lead to overfitting if too many parameters are included in the fit. To test this possibility, denss.pdb2mrc.py was rerun on all eight datasets where the option to fit *ρ*_0_ and *δρ* was disabled. For comparison, CRYSOL was also rerun while disabling minimization of its fitting parameters. The results in Table 2 (columns labeled “No Fit”) show a significant increase in *χ*^2^ relative to the fitting results shown in Table 1 (ranging from 4% to 550% increase, median of 36%). This demonstrates that fitting *ρ*_0_ and *δρ* does indeed have a measurable and significant effect on the quality of the fits produced by denss.pdb2mrc.py, despite the values changing only slightly during fitting. However, these results also show that denss.pdb2mrc.py is able to produce significantly better fits than CRYSOL without optimization of parameters (which ranged from 41% to 2650% increase, median of 661% increase). This suggest that the approach outlined above for estimating the excluded volume is accurate, resulting in not only the removal of a free fitting parameter, but also a significant increase in the accuracy of the remaining parameters. Taken together, these results demonstrate a significant improvement of the parameter-free calculation of scattering profiles over previous methods. However, more accurate estimations of the bulk solvent density and the hydration shell will be required to fully remove the need for fitting, which may be provided by more advanced algorithms invoking MD or other explicit solvent modeling.

**Table 2.**
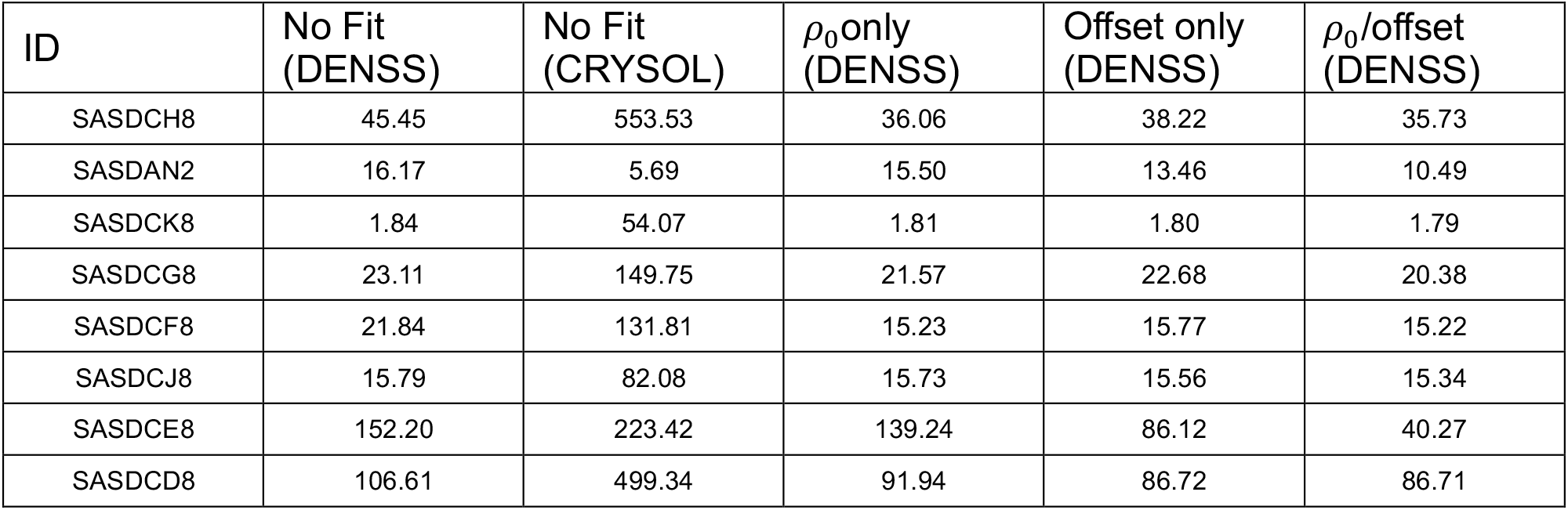
Comparison of the effect of disabling fitting parameters on χ^2^. All fits have been performed with the shell contrast fitting disabled.

The fitting procedure described above uses least squares to determine the linear scale factor that scales the data to the calculated profile. There also exists an option to fit an additional constant offset using least squares which is disabled by default. This option aims to correct for possible background subtraction errors in the data and is thus added directly to the data (rather than subtracted from the calculated profile). To test the effect of fitting this constant offset, all eight datasets were rerun using denss.pdb2mrc.py while enabling this option (while disabling fitting the hydration shell contrast). The results are shown in Table 2. In each case, the *χ*^2^ value was seen to improve by a small margin compared to fitting only *ρ*_0_. However, when only fitting the offset, the improvement in *χ*^2^ over no fitting was seen to be very similar to fitting only *ρ*_0_. In other words, fitting either the bulk solvent density or the constant offset results in a very similar effect on the *χ*^2^. This suggests fitting the constant offset by itself may have a detrimental effect on the experimental data, and thus this option is disabled by default.

In each case above, the option to include atomic B-factors as described by Equation 6 was disabled. To assess the effect of modeling crystallographic B-factors provided by the atomic models, denss.pdb2mrc.py was rerun on all eight datasets where the B-factor calculation was enabled. In all eight cases inclusion of the B-factor significantly reduced the quality of the fit to the experimental data, where *χ*^2^ typically increased by a factor of 2 or more (not shown). Due to this result, the default behavior for denss.pdb2mrc.py is to disable inclusion of atomic B-factors. It is possible that this effect may be due to using crystallographic B-factors where the crystal has been cryogenically frozen in contrast to the room temperature solution scattering experiment, thus resulting in inaccurate B-factor estimates for the SWAXS experiment. However, given that the increase in temperature in the SWAXS experiment will only increase the thermal motion of the atoms, it is unlikely that room temperature B-factors would improve (and given the results above would likely worsen) the quality of the fit to the data. To verify this, a room temperature structure of lysozyme (PDB ID: 5NJM (50)) was fit to the SASDCK8 experimental data using room temperature B-factors. Indeed, this resulted in a further increase in *χ*^2^ for the room temperature model (*χ*^2^ = 13.05) relative to the model with cryogenic B-factors included (*χ*^2^ = 2.51), both of which are higher than using either model without B-factors (*χ*^2^ ≈ 1.75 in both cases) (Fig. 6).

**Figure 6.**
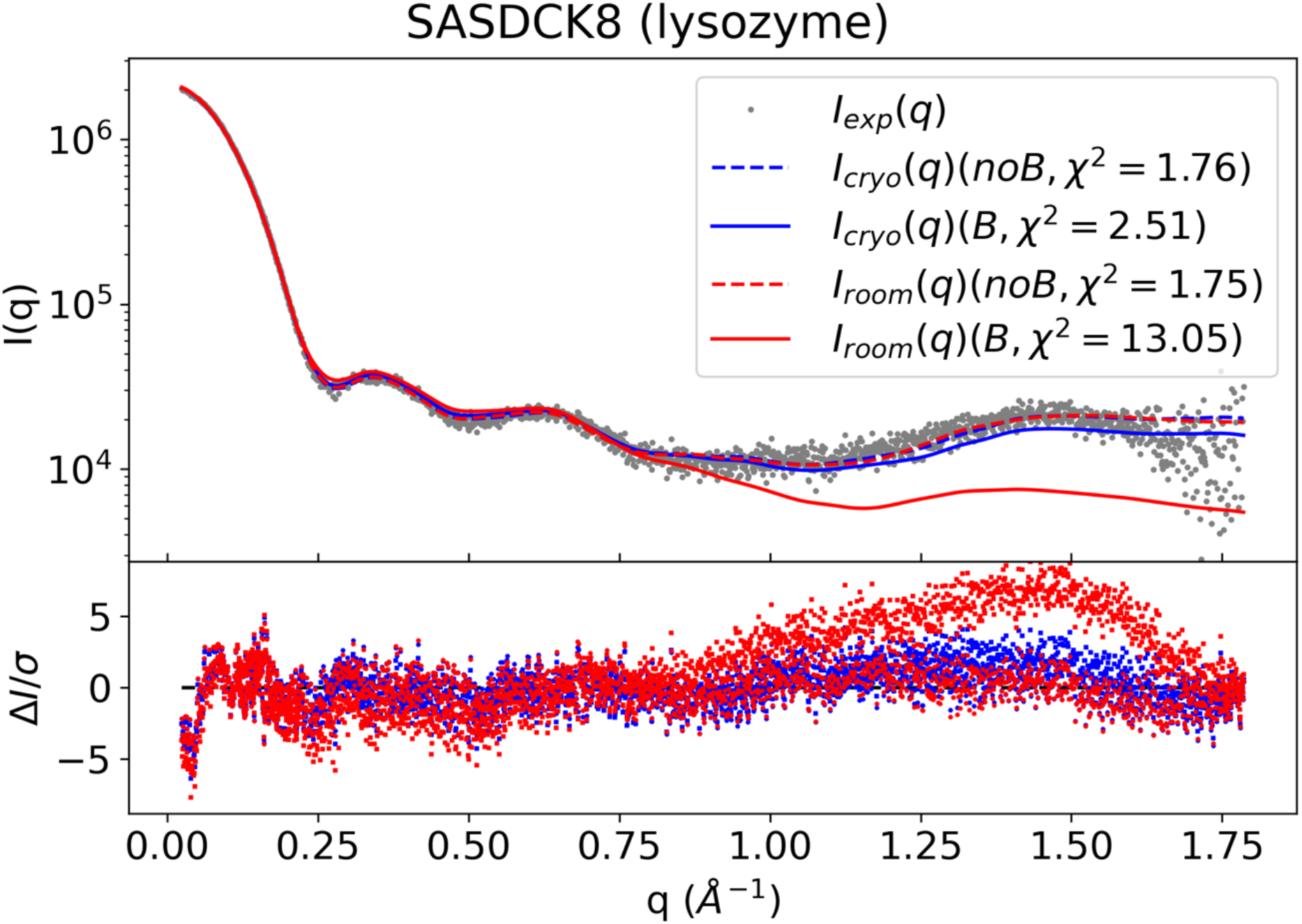
Comparison of the effect of B-factor on calculated scattering profiles. Experimental scattering data from SASDCK8 are shows as gray dots. Blue curves represent the scattering calculated from the atomic model determined under cryogenic conditions (SASDCK8 atomic model), and red curves represent the atomic model determined under room temperature conditions (PDB 5JNM). Dashed lines correspond to the default calculation of denss.pdb2mrc.py with B-factors ignored and solid lines correspond to the denss.pdb2mrc.py calculation with B-factors included. χ^2^ values are shown in the legend for each calculated profile. Normalized residuals are shown at the bottom.

The excluded volume estimation uses a sum of Gaussians approximation to model the bulk solvent density as dummy atoms, as is commonly done in algorithms such as CRYSOL and FoXS. Other algorithms have sought to model the excluded volume as a flat density by modeling it as a collection of homogenous cubes as described above. The modeling of density on a uniformly spaced cubic grid in denss.pdb2mrc.py makes it fairly trivial to model the excluded volume density using this cube method by simply setting the voxels within the solute atoms’ van der Waals radii to the bulk solvent density. This approach is similar to the “union of spheres” method implemented in SoftWAXS (18). To compare the default method of modeling the excluded volume as a sum of Gaussians with the flat bulk solvent cube method, all eight datasets were rerun using the flat cube method. In each of the eight cases, the fit of the atomic model to the experimental scattering profile was significantly worse when using the flat cube method compared to the sum of Gaussians, with *χ*^2^ increasing by a factor of 3 on average (not shown). Given these results, the default option in denss.pdb2mrc.py is set to use the sum of Gaussians approach.

While the goal of the present study is the generation of an accurate density map fitting experimental SWAXS data and not computational speed, our approach inherently results in a favorable scaling relationship relative to the number of atoms, as shown in Table 3. Calculations utilizing the full Debye equation scale quadratically with the number of atoms. However, calculating density in real space followed by an FFT enables efficient calculation of Fourier terms, since the real space density calculation can be localized to a small number of voxels in the vicinity of each atom. In each case tested, the computational speed is favorable to CRYSOL, which implements the fast spherical harmonics approximation, while producing similar quality fits. To further test the speed for particularly large macromolecules, additional models were downloaded from the PDB containing more than 100,000 atoms up to ∼2 million atoms (Table 3). As these molecules lack any publicly available experimental scattering profiles, no fitting was performed for either denss.pdb2mrc.py or CRYSOL. Additionally, to evaluate the effect of the fitting procedure on execution time, scattering profiles were recalculated without fitting for the models in Table 1, as well. The execution time when fitting using denss.pdb2mrc.py for the models in Table 1 ranged from 2.06 s to 34.49 s, and without fitting ranged from 1.23 s to 16.71 s. On average approximately half of the total execution time was taken up by fitting. This is in contrast to CRYSOL, which showed nearly identical times (within 1% typically) regardless of fitting, ranging from 5.43 s to 109.63 s. The larger relative fitting time for denss.pdb2mrc.py may result from using the Nelder-Mead simplex algorithm for numerical optimization, rather than performing a grid search as done in CRYSOL. This suggests that there may be room for improving the computational efficiency of denss.pdb2mrc.py in the fitting step. Even with fitting, denss.pdb2mrc.py ranges from about two to twelve-fold faster than CRYSOL in each case. When considering larger macromolecules (>100,000 atoms) without fitting, denss.pdb2mrc.py is on average ten-fold faster than CRYSOL, with larger macromolecules experiencing greater gains (Table 3).

**Table 3.** Times of execution for denss.pdb2mrc.py (“DENSS”) and CRYSOL. The column labeled “ID” shows either the SASBDB ID or the PDB ID. Times are given in seconds, calculated using the zsh “time” command.

| ID | N<sub>atoms</sub> | $t_{DENSS}$ (s) (fit) | $t_{CRY SOL}$ (s) (fit) | $t_{DENSS}$ (s) (no fit) | $t_{CRY SOL}$ (s) (no fit) |
| --- | --- | --- | --- | --- | --- |
| SASDCH8 | 1744 | 2.06 | 24.88 | 1.23 | 20.63 |
| SASDAN2 | 1852 | 3.07 | 5.43 | 2.00 | 5.45 |
| SASDCK8 | 1959 | 3.36 | 5.74 | 1.63 | 5.74 |
| SASDCG8 | 4035 | 4.93 | 23.45 | 2.39 | 23.88 |
| SASDCF8 | 8387 | 17.14 | 30.53 | 8.00 | 30.85 |
| SASDCJ8 | 23848 | 19.40 | 56.48 | 10.18 | 57.35 |
| SASDCE8 | 30708 | 34.49 | 67.98 | 10.92 | 68.60 |
| SASDCD8 | 66456 | 31.44 | 109.63 | 16.71 | 110.70 |
| 1EI7 | 86450 | - | - | 23.31 | 172.55 |
| 1R27 | 121545 | - | - | 29.01 | 278.13 |
| 1T0T | 131235 | - | - | 24.76 | 267.79 |
| 1A6C | 240960 | - | - | 44.91 | 484.80 |
| 7OJ7 | 393780 | - | - | 78.36 | 815.06 |
| 1OHF | 1057435 | - | - | 168.24 | 2165.00 |
| 3J3Q | 2440800 | - | - | 362.45 | 4657.07 |

## 4. Discussion

Here we have presented our approach to solving the difficult problem of predicting accurate solution X-ray scattering profiles from atomic models at wide angles. Our approach generates an electron density map at high resolution, which may be useful for subsequent modeling calculations. Unique to our method is the calculation of atomic volumes directly from the atomic coordinates for each atom for the calculation of the excluded volume density. By calculating unique atomic volumes, we have removed one of the most significant fitting parameters commonly employed in popular algorithms such as CRYSOL and FoXS, namely the global expansion factor which modifies the radius of each atomic group. Despite removing this fitting parameter, our approach yields highly accurate fits to all eight experimental SWAXS datasets tested. Additionally, the observation that the two fitting parameters (namely the bulk solvent density and the hydration shell contrast) displayed narrow distributions near the default values suggests that the method is robust to overfitting. It may be possible that coupled with accurate predictive calculations of the bulk solvent density and the hydration shell density that no fitting parameters will be necessary in the future. Such a development would be very powerful for the community, enabling significant improvements in the accuracy of modeling algorithms while reducing the risk of overfitting. As SWAXS contains limited information compared to other structural methods, this would be a significant development in the field.

Our results above showed that fitting a constant offset has a similar effect on the *χ*^2^ value as fitting the bulk solvent density. It is important to note that the subtraction of a constant offset from the experimental data is a fairly common approach to deal with background subtraction errors (the option is available in CRYSOL (16), though disabled by default, and implemented in some MD SWAXS algorithms (21)). Most MD-SWAXS algorithms do not fit the bulk solvent density in order to avoid unnecessary fitting parameters. However, our results show that fitting the offset has a similar effect on *χ*^2^ as fitting the bulk solvent density, and thus fitting the offset may be masking additional information in the SWAXS profile, in particular at high q values. Indeed, this has been suggested before (21), and the only solution currently available to truly avoid such overfitting is to fit neither parameter in the MD simulations and to simply require highly accurate background subtraction experimentally. Additionally, it may be possible to use the knowledge of the components in the solution (e.g., salt concentrations, etc.) to estimate the bulk solvent density by direct calculation, though it is unknown if this approach is accurate enough in practice. As described above, our approach results in a very narrow distribution of bulk solvent density values even without fitting a constant offset, suggesting this approach is highly accurate and robust to such overfitting concerns.

Several important findings were observed in our results. The incorporation of crystallographic B-factors (both at cryogenic and room temperature) significantly decreased the quality of the fits to the experimental data. These results suggest the method for incorporating B-factors in solution scattering is different than in crystallography (51). In crystallography, the atomic form factors of the atoms are directly modified, and thus the contribution of the B-factor is additive in terms of structure factors. However, in solution, since particles are assumed to be dilute and non-interacting, the B-factor contributions should add only in intensities, not structure factors. While the incorporation of isotropic B-factors in Equation 6 derived from crystallography has been used in other software for calculating SWAXS profiles (17,22), this approach clearly requires modification to properly account for thermal disorder in a solution experiment. One such modification has been previously derived mathematically by Moore (51), which outlines a first order approximation for this effect using the Debye calculation. Such a modification cannot be directly included in our approach, as we do not use the Debye equation. However, it is possible that an alternative derivation, or direct modeling of an ensemble, may be possible in future versions. Such a calculation would require an ensemble of states, where the intensities of each member of the distribution of states dictated by the B-factor would be calculated and added. Future improvements to denss.pdb2mrc.py may allow for modeling this effect accurately.

We employed the common approach of modeling the excluded volume as a sum of Gaussian dummy atoms centered at the protein atom coordinates. Comparison of this approach with the cube method (28), where the bulk solvent is modeled as a flat value subtracted from each protein atom grid point, showed that in our algorithm the sum of Gaussians approach fits the data better than the cube method. This finding is in contrast to other algorithms such as SoftWAXS which showed the opposite effect (18). However, it is important to note that in our approach, we do not fit the excluded volume radii independently for each protein, in contrast to the method implemented in SoftWAXS. As described above, the excluded volume radii were fit once and are held constant for all subsequent samples. This difference in approach may explain why the cube method in our implementation results in a worse fit, as the radii have not been optimized separately for each protein.

Our method was also shown to be computationally efficient, favorable to the leading CRYSOL software. The primary reason for the computational efficiency of our approach is the logic of using real space density prior to calculation of reciprocal space structure factors. Since density decays rapidly as a function of distance from the center of an atom, the calculation of density can be limited to only a handful of voxels near the atom. This is in contrast to methods that directly calculate Fourier terms from atomic coordinates since the Fourier terms must be calculated at all sampled locations in reciprocal space. While our method does require the additional step of calculating the Fourier transform of the density map, this calculation is very fast using FFT algorithms in Python (currently using either NumPy or SciPy (10,52,53)). Additionally, this portion of the calculation does not scale with the number of atoms, but instead scales with the number of voxels. The scaling relationship is highly favorable as well, given the efficiency gains implemented in modern FFT algorithms that scale as O(n log n) (10). As our aim was not explicitly to create a fast algorithm, it is expected that optimizations to the code may significantly improve calculation time. Such improvements may include using more highly optimized FFT algorithms such as FFTW (54) or GPU acceleration (55-57). Additionally, since the real space calculations of density are a simple sum over all atoms for both the *in vacuo* and excluded volume terms, these calculations are likely to be highly amenable to parallel computing. Similarly, the distance transform calculation for the hydration shell may also be improved using parallel computing. If such efficiency gains to the calculation of the scattering profile are realized in future versions of our algorithm, the approach could be implemented in atomic modeling programs such as MD that use experimental SWAXS data to guide the simulation, which require hundreds of thousands of scattering profile calculations. Coupled with the reduced risk of overfitting as described above, these advances may enable a significant improvement in the accuracy and utility of SWAXS-driven MD algorithms to model structural dynamics of proteins in solution (35,58).

Finally, the algorithm presented here generates a highly accurate electron density map of a protein at high resolution that agrees with experimental SWAXS data. This will enable future modeling applications using our novel iterative structure factor retrieval algorithm implemented in DENSS (9). By combining experimental SWAXS data of full-length proteins or protein-protein complexes with high resolution density maps from known structural fragments or subunits or using predicted structures (e.g., from AlphaFold (59)), we will be able to significantly improve the resolution and accuracy of density reconstructions of unknown regions. This promises to improve the ability to interpret structure and dynamics of proteins and other biomacromolecules in solution states using SWAXS.

## 5. Author Contributions

TDG conceived and designed the project. SM assisted with the initial implementation of the algorithm. TDG and SRC wrote the code, analyzed the data, and wrote the manuscript.

## 7. Acknowledgements

The authors would like to acknowledge funding from the National Institutes of Health National Institute of General Medical Sciences through award R01GM133998 and from the National Science Foundation through the BioXFEL STC 1231306.

